# Height differences in clonal stands of *Tussilago farfara* promote outcrossing by influencing pollinator behaviour

**DOI:** 10.1101/2020.02.17.952713

**Authors:** Aleksandra J. Dolezal

**Affiliations:** Department of Integrative Biology, University of Guelph, Guelph, Ontario, Canada

**Keywords:** Insect behaviour, outcrossing, mixed mating system, plant height, *Tussilago farfara*

## Abstract

Plants with mixed mating systems balance the advantages of selfing and the costs of inbreeding. Previous studies have shown that plant species with the ability to self-pollinate and cross pollinate have strategies to promote outcrossing to increase genetic diversity. Various features of floral morphology are thought to be deliberate mechanisms to manipulate plant mating systems. I hypothesized that within-plant variation in flower stem height is a morphological trait that can reduce geitonogamy by increasing pollinator movement among plants. This hypothesis was tested using coltsfoot (*Tussilago farfara*); coltsfoot plants host several flowering stems that differ in height, with each stem having a single, terminal flowerhead. I used cut flowers to create ‘populations’ of coltsfoot in which each plant had four inflorescences with equal or unequal flower stalk lengths and measured frequency of insect pollinators that would stay among flowers within a plant or move to other plants. All pollinators (bee flies, hover flies, solitary bees and wasps) showed a marked discrimination in favor of leaving plants with flowers of different heights and stayed when plants had flowers of the same height. This study shows that variation in flower heights is important for reducing geitonogamy in coltsfoot and suggests that the evolution of this morphological trait should be considered in studies of mating systems.

## Introduction

One of the fundamental questions in ecology is how plant morphology affects pollinator behavior (Chaffey, 2014). In particular, how does plant morphology indirectly affect reproductive strategy when a species can both self-pollinate and cross-pollinate? Many plants in nature employ mixed mating strategies to better increase their chances of becoming fertilized and to increase seed set (Pannell et al., 2013). However, a reproductive strategy to promote outcrossing is of critical importance for the adaptation and evolution of all eukaryotic organisms (Raven et al., 2005).

Plants have methods to reduce or avoid geitonogamy in order to increase the amount of genetic variation within and among populations (Loveless and Hamrick, 1984). Separate male and female flowers may both occur on the same plant or occur at separate times when they each reach maturity during different periods in monoecious plants, however some plants produce staminate and pistilate flowers on separate plants (Raven et al., 2005). This strategy occurs in dioecious plants where flowers are constructed such that stamens and stigmas never meet each other to rely exclusively on outcrossing. In this extreme case, pollen must be transferred to the stigma of another flower rather than itself which never allows self-pollination to occur.

In recent years, studies have looked at the relative importance of plant traits on pollinator visitation rate and the role that these floral morphological traits have on the fitness of the plant species. If a plant is multiflowered, this may be important to attract pollinators by having a big enough display, but there is a risk that a flower visitor will go from one flower to another on the same individual, thus increasing geitonogamy (Wilmer, 2011). Understanding how foraging choices occur at the local scale is essential because these choices may directly affect plant fitness (Lazaro and Totland, 2010). Many different processes are involved in explaining plant morphological traits into realized success for insect pollinated plants, such as the effects of plant traits directly on the behavior of insect pollinators (Mitchell, 1994). For example, pollinator visitation rates differ among plant species due to the response to traits such as floral size, floral shape, color preference, and the rate of nectar production (Mitchell and Waser, 1992; Campbell, 1991). However, little is known about local foraging decisions for the majority of pollinator guilds, with the exception of bumblebees (Stout et al., 1998).

Several studies have investigated the effects of plant morphological traits in manipulating pollinator behavior and mating cost in hermaphroditic species. For example, Harder and Barrett (1995) found that large floral displays of *Eichhornia paniculata* had enhanced pollinator attraction, however endured higher selfing and lower outcrossing rates due to pollen discounting by bees. Similarly, Williams et al. (2001) investigated a different morphological trait, the number of inflorescences per plant, and showed that distinct plants with multiple inflorescences attract more pollinators than those with single inflorescences. In the context of plant fitness, increased visitation to plants with multiple inflorescences may also reduce the fraction of selfed seeds (Miyake and Sakai, 2005), due to the greater import of pollen from other plants. Overall, this research highlights that a number of plant morphological traits influence pollinator behavior and affects mating costs in self-compatible species.

Although studies have illustrated how several morphological traits alter pollinator movements and outcrossing rates, some traits, such as flower height, remain less understood. Previous studies of selection for floral traits indicate plant height as an important trait for pollination success (Johnston 1991). Lortie and Aarssen (1999) found that taller plants of the biennial weed *Verbascum thapsus* attracted more pollinators, showing that they may experience higher rates of outcrossing and have more diverse pollen donor arrays. Hingston and Potts (2005) found that the gum tree *Eucalyptus globulus* had higher outcrossing rates on its upper branches due to the flowers occurring at different heights. This resulted in birds foraging more in the upper canopy region. Height variation of flowers may attract different types of visitors, or different visitation rates of these individual visitors (Wilmer, 2011). Many species of plants present flowers at multiple heights, in some cases this variation in flower height may be due to architectural constraints of the plant, in others, however, there is no clear reason for this variation in flower height. For example, coltsfoot (*Tussilago farfara*, Asteraceae) has terminal flowerheads on stalks that emerge from the soil and differ in height. Nonetheless, these stalks typically grow longer and more uniform post-fertilization and prior to seed release, suggesting that resources or architectural constraints do not drive intraspecific variation (Bakker, 1960). As a result, it is unclear why this height variation and height constancy persists in this species.

I hypothesized that differences in flower heights within a plant would increase pollinator movement among plants, thus decreasing geitonogamy in *T.farfara.* To test this, I introduced two different treatments to pollinators –treatments with *T. farfara* flowers at different heights and treatments with *T. farfara* flowers at the same height. This distinguishing character enabled me to quantify pollinator behavior in different height treatments. I assessed the number and diversity of pollinator arrivals to treatments, and whether pollinators decided to stay within clonal stands or leave clonal stands in each treatment. To my knowledge, this is the first study to determine the effect of different flower heights on pollinator preference and geitonogamy.

To examine the effect of flower height arrangement on pollinator behaviors, I addressed the following questions in my study: (i) Will height variation affect the number of pollinator visitations? (ii) Will the presence of flower height differences encourage pollinators to leave clonal stands? (iii) Do different types of *T. farfara* flower visitors (bee flies, hover flies, solitary bees, and wasps) respond differently to within-plant variation in flower height?

## Methods

### Study species

*Tussilago farfara* is a small, clonal, herbaceous perennial in the Asteraceae family. It has been introduced to North America and has a wide distribution throughout the continent (Myerscough and Whitehead, 1996). It is commonly found in clusters of multiple height stems (5 cm - 30 cm) produced by a single rosette, with hermaphroditic flowers. The flowers appear early in spring, prior to the emergence of any leaves when increasing temperatures initiate elongation of the inflorescence stems followed by flowering (Balchin and Pye, 1950). The golden-yellow flower heads contain an outer region of numerous female ray florets and inner region of 30-40 male disc florets (Pfeiffer et al., 2008). The flower heads are protogynous, in which the female reproductive organs mature before the male reproductive organs (Myerscough and Whitehead, 1996). This species has a mixed mating system, and can both self-pollinate and outcross. A study on the reproductive strategy of *Tussilago farfara* by Ogden (1974) found that, in dense populations, vegetative reproduction fails, and seed production suffers little. Reproductive tactics tend to favor seed production relative to clonal spread. At the study site, it is mainly visited by hover flies, bee flies, and solitary bees, but wasps were observed and also included in this study.

### Study area

Field work for this study was conducted from May 10 to May 13, 2016 with a population of *T. farfara* located at Koffler Science Reserve at Jokers Hill (King township, Ont.; 44°03N, 79°53’W). Samples were taken from three populations located within 1 km of each other. All sites were in partial to full sun and were along roadsides or trail sides in areas that are regularly disturbed by mowing.

### Experimental design

I designed two treatments to determine whether variation in stem height would alter pollinator behavior. The two treatments consisted of a different height treatment with stem heights consisting of 6 cm, 10 cm, 14 cm, and 18 cm and a same height treatment consisting of four flowers all 12 cm. These heights were chosen to reflect natural variation in flower stem length, the mean length of *T. farfara* in the study sites being 12 cm. Flowers were cut at the start of the day when they were open, and flowers of a similar size and age were chosen to make sure there was no bias from pollinators to visit stands of different flowers.

In each experimental unit, I cut and manipulated stems of four flowers to be placed into a vial. Each of these experimental units consisted of four flowers in a vial with soil from the study site and water to keep plants from wilting during observations. Two experimental units were placed 30 cm apart (the average distance observed in natural populations) from each other in a single block, and treatments within a block contained two replicates of each treatment (Figure 1). These different treatment combinations were used to ensure pollinator behavior is not influenced by nearby flowers.

**Figure 1:**
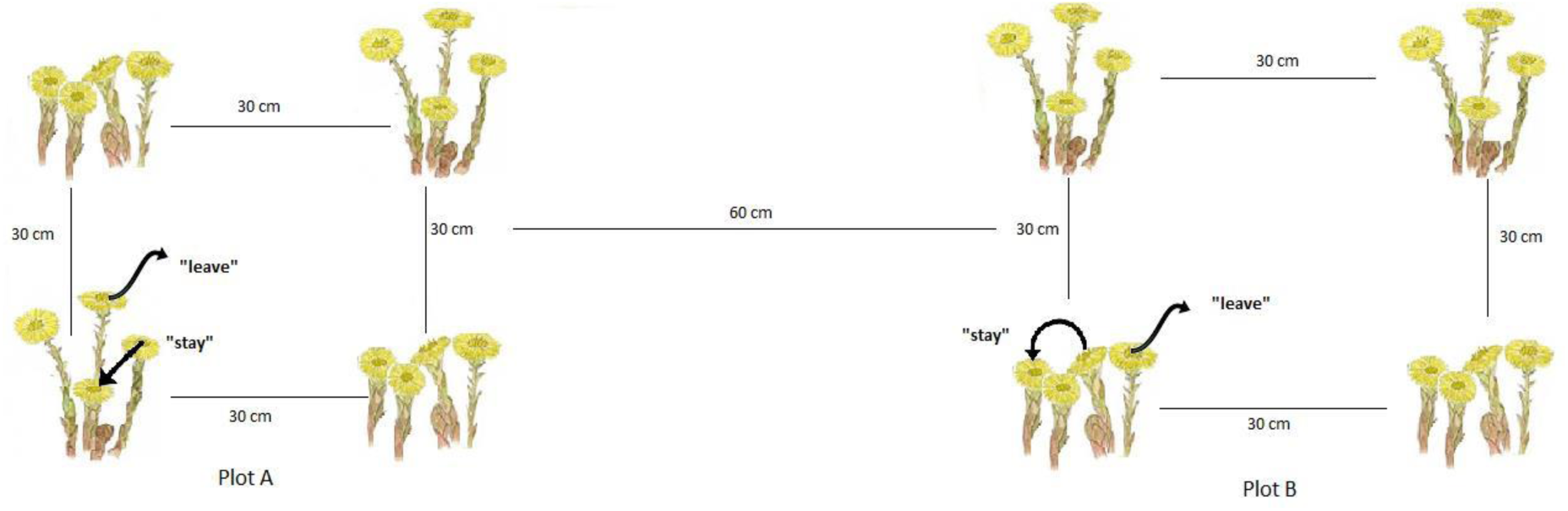
Experimental design consisting of two treatments: different height (DH) and same height (SH). Plot A and Plot B include a combination of two DH and two SH treatments, A has vials diagonally across from each other and B has treatments parallel to each other. A pollinator’s action was considered “leave” if it landed on a flower and its next decision was to fly away or fly to another clonal stand. An action was considered “stay” if a pollinator’s next decision was to land on any of the four flowers within the clonal stand.

### Data collection

During May 11 to May 13, plants were observed for insect pollinators. All observations were conducted in sunny stands starting at 11:30 am – 1:30 am unless wind or rain disrupted foraging behavior, then different times of the day were chosen to accommodate optimal sun exposure. I observed each treatment plot for insect pollinators for two hours. Pollinators were observed approximately 2 m away from plots using binoculars. Pollinators did not appear to alter foraging behavior in response to my proximity. A visit was defined to have occurred when the visitor’s body contacted the reproductive organs (ray or disc florets) of the flower. The insects’ next decision was observed and recorded; moving to another flower in the same vial was recorded as ‘staying’, whereas moving to another flower in a different vial was recorded as ‘leaving’. Once a pollinator was recorded as staying or leaving, no further observations were made for that pollinator during that visit (because the individual pollinators were not marked, repeat visits by individual pollinators over several hours or days may have occurred). I categorized visitors to one of four groups: bee flies, hover flies, solitary bees and wasps. Each plot was located near a stand of *T. farfara* to get the same pollinator groups. *T. farfara* was chosen because they are common at Koffler Scientific Reserve and were in bloom early May. No other plant species bloomed in the study areas except for dandelions which have similar pollinators as the study species.

### Statistical analysis

I used a Pearson’s Chi-squared test with Yates’ continuity correction via RStudio (RStudio, 2015), to examine the effects of plant height on pollinator visitation behavior. The response variable was the leave/stay (0 / 1) action observed in each treatment. In this analysis, data from all three sites were pooled to increase sample size and because the results did not differ between sites (results not shown). Fishers Exact Test was used to obtain p-values for pooled samples.

## Results

A total of 190 pollinators were observed May 11 to May 13, 2015. The relative abundance of four pollinator types varied across populations (Figure 3) and both solitary bees and hover flies were the dominant visitors to *T. farfara* stands. Neither treatment nor any interaction effects of treatments had an effect on insect visitation rates – there was no significant difference in the frequency of pollinators that visited the ‘different height’ treatment (94/190) compared to the ‘same height’ treatment (96/190; Figure 2, x^2^=0.0046, DF=1, P=0.9458). These results indicate that there was no preference for one type of treatment.

**Figure 2:**
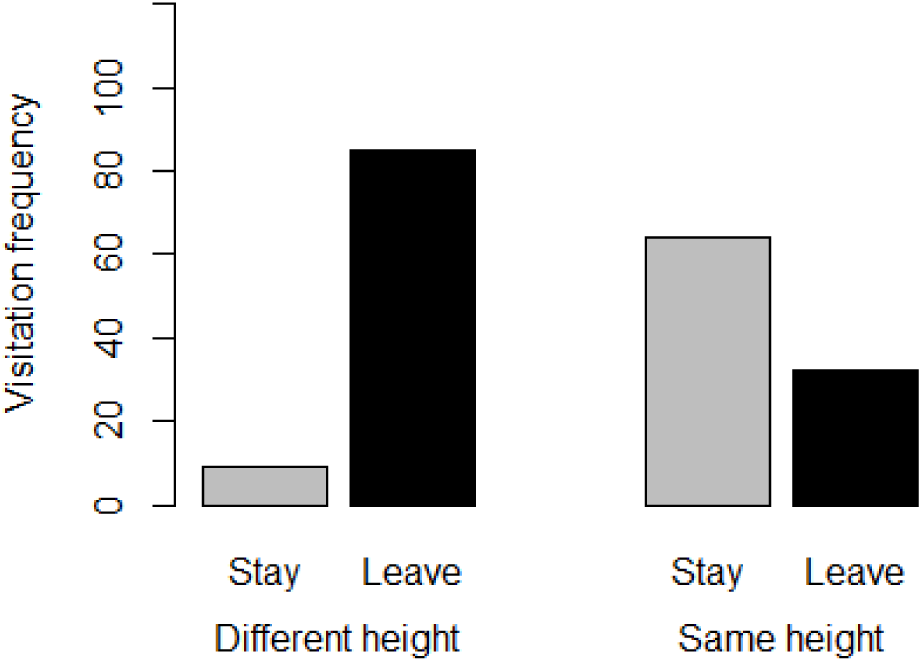
Frequency of pollinator visitations to *T. farfara* occurring in plots with different treatments in three study populations (n(different height)= 94, n(same height)=96)).

**Figure 3:**
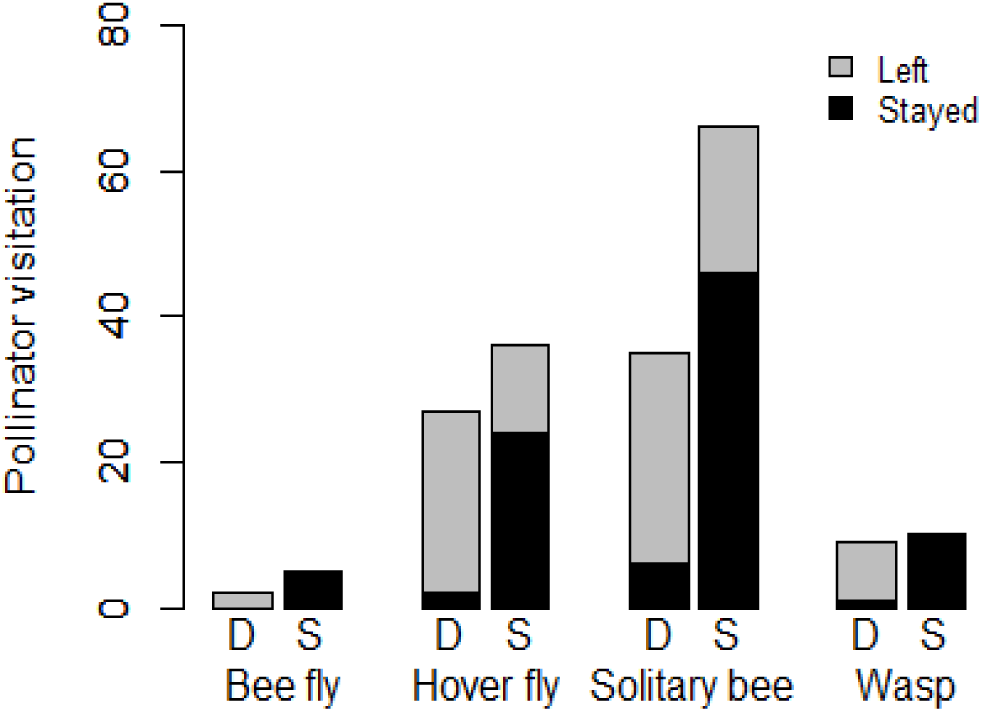
Pollinator visitation behavior occurring in height treatments of *T. farfara* plots. P- values are based on Fisher’s Exact Test. No difference between pollinator behaviors in treatments (D represents different height treatment and S represents same height treatment). Grey and black colors represent the action of leaving a stand or staying within a stand, respectively.

The height difference treatment has a significant effect on pollinator behavior. When inflorescences were presented at different height, insects were more prone to leave a stand of flowers, whereas they were more likely to stay and forage within a stand when inflorescences were presented at the same height (Figure 2; x^2^=63.042, DF=1, p<0.001). This behavior of pollinators suggests that geitonogamy will be reduced in *T.farfara* when inflorescences differ in height, as pollinators are more likely to visit flowers of other plants.

Pollinators showed no difference in behavior in treatments (Figure 3). All showed the general trend of leaving treatments of different height flowers and staying within treatments of same height flowers. The bee fly showed the most significant trend, in which it always left a different height stand and always stayed within a same height stand (N=7, p<0.05). Other pollinators include: hover fly (N= 63, p<0.001), solitary bee (N=101, p<0.001) and wasp (N=19, p<0.001).

## Discussion

In this study, I have demonstrated that height variation affects pollinator behavior and consequent reproductive strategy in populations of *T. farfara.* Generally, pollinators that landed within a different height treatment left the clonal stand and those that landed on a same height treatment stayed within a clonal stand. My data suggests that even though there are a variety of pollinators visiting flowers (Figure 3), they all exhibit the same behavior: leaving different height flowers and staying within same height flowers (Figure 3). My findings support the hypothesis that hermaphroditic plant morphology in *T. farfara* promotes outcrossing with the use of height variation to manipulate insect pollinators. This study is the first to document that natural height variation in *T. farfara* influences pollinator behavior and promotes outcrossing. Due to the fact that my results are based on observations of flowers under natural conditions, they could potentially support the association of flower stem height variation and reduced geitonogamy in other taxa of the Asteraceae family.

Despite the striking results of my analyses, it is worth considering the limitations imposed by my experimental design and study overall. First, my study was observational and there may have been misidentification of insect pollinators that visited plots as some flew away too quickly to get a sure identification through binoculars. Due to the close proximity of the 4×4 grid treatments there may have been times when I was viewing one stand and pollinators came into the other that I did not record. For a future study, perhaps another observer could assist me in recording data. Second, two days of observations were cloudy and had 45 km/h wind that offered a limited window to observe pollinators. Pollinator flight activity is constrained by severe weather conditions (Totland and Matthews, 1998) such as wind speed or temperature which could influence pollinators to forage differently. Third, depletion of nectar resource from disc rays of the flower could have influenced more pollinators to leave clonal stands if they experienced a no-nectar flower within that stand. Studies have suggested that a pollinator (specifically bees) leaves a plant if it comes across one or two nectar-poor flowers (Kadmon and Shmida, 1992). Studies of bumble bees determine that they visit other flowers on individual inflorescences if those flowers provide nectar reward (Cresswell, 1990; Johnson et al., 2004) and if the costs of foraging for previously unvisited flowers remain fairly small (Ishii et al., 2008). A pollinator should leave an unrewarding plant as soon as possible but stay and continue to visit the flowers on a rewarding plant. It would be worthwhile to test the pollinators of *T. farfara* to see if they show similar behaviors in foraging when nectar is added to flowers. Fourth, the flower orientation could have influenced pollinator behavior in leaving or staying within stands. The different height treatment flowers were at angles because of the weight of the taller stems inching forward, while the same height flowers stood upright in the same horizontal positions. Floral orientation is thought to affect pollinator attraction, foraging behavior, and pollen transfer (Wang et al., 2014). However, previous studies are on species of plants that produce single flowers, and rather little is known about the effect of floral orientation on pollination in more multifaceted inflorescences. Future research is necessary to examine the effect of flower orientation on pollen import and export, as well as the visitation behavior of pollinators, in order to understand the influences that horizontal orientation or vertical orientation may have on outcrossing success.

Future research should investigate whether this pollinator behavior indeed increases plant fitness and how flower stem height affects probing times. Carromro and Hamrick (2005) in their study of *Verbascum thapsus* demonstrated that tall plants receive larger and more genetically diverse pollen loads, while shorter plants experience pollen limitation which lead to increased selfing. However, similar to my study, the authors did not look at times spent on flowers of different heights which could also affect plant fitness. During my observations I noticed that pollinators seemed to probe disc rays on the same height treatment for longer than on the different height treatments. It would be beneficial to add duration of probing time as another variable in my study to determine how pollinator behavior may be affected. Since pollinators can see that flowers are close by, I would predict they would have fewer probing rates on same height treatments. This is because the costs of searching is not constrained for flowers side by side. Waddington and Holden (1979) proposed that honeybee visitation to flowers is based on an optimal foraging model where a bee achieves maximum caloric intake by choosing to visit the closest flower. It would be worthwhile to see if seed set is higher for treatments with longer probing times, as this would increase individual plant fitness.

A study by Waddington (1979) states that there is greater selection against plants of heights where the probability of interspecific pollinator flights is at its highest, and therefore plant species partition pollinators by presenting inflorescences at different heights. He suggested that pollinators shared by the species exhibited the same horizontal flight pattern, where they tended to remain at a certain height when foraging between flowers. Similarly, I have shown that plants presenting flower heads at different heights would reduce geitonogamy because pollinators would be more likely to move horizontally rather than vertically. Plant height differences can have a direct effect on pollinator behavior and thus, resulting in an indirect effect on reproduction. Considering the mating costs of geitonogamy, selection should present floral traits that reduce geitonogamy in clonal plants with a clumped architecture, such as limiting same height inflorescences (De Jong et al., 1992). The magnitude of the potential fitness consequences associated with geitonogamy in *T. farfara* and how they are balanced by the reproductive benefits of flower height display are currently unknown. Species with mixed mating strategies present unique opportunities for understanding the evolution of mating systems. In the case of *T.farfara*, it is further complicated by the indirect effect that inflorescent height has on promoting outcrossing rates by manipulating foraging decisions in insect pollinators.

## Acknowledgements

I thank J. Eckenwalder and B. Gilbert for valuable comments on early versions of this experiment and in all aspects of this work. I thank A. Korda for assistance in the field and in harvesting flower stems. I thank S. Wadgymar with assistance in statistical analyses.

